# Impaired extinction of cocaine seeking in HIV-infected mice is accompanied by peripheral and central immune dysregulation

**DOI:** 10.1101/2023.08.11.552858

**Authors:** Lauren A Buck, Qiaowei Xie, Michelle Willis, Christine M Side, Laura L Giacometti, Peter J Gaskill, Kyewon Park, Farida Shaheen, Lili Guo, Santhi Gorantla, Jacqueline M Barker

## Abstract

Substance use disorders (SUDs) are highly comorbid with HIV infection, necessitating an understanding of the interactive effects of drug exposure and HIV. The relationship between progressive HIV infection and cocaine use disorder is likely bidirectional, with cocaine use having direct effects on immune function while HIV infection can alter addiction-related behavior. To better characterized the neurobehavioral and immune consequences of HIV infection and cocaine exposure, this study utilized a humanized mouse model to investigate the outcomes of progressive HIV infection on cocaine-related behaviors in a cocaine conditioned place preference (CPP) model, and the interactive effects of cocaine and HIV infection on peripheral and central nervous system inflammation. HIV infection did not impact the formation of a cocaine CPP, but did result in resistance to extinction of the CPP. No effects of HIV on yohimbine-primed reinstatement or cocaine seeking under conflict were observed. These behavioral alterations were accompanied by immune changes in HIV infected mice, including increased prefrontal cortex astrocyte immunoreactivity and brain-region specific effects on microglia number and reactivity. Peripheral immune system changes were observed in both mouse and human markers. Among other targets, this included HIV-induced reductions in mouse IL-1α and G-CSF and human TNFα and cocaine-induced alterations in human TNFα and mouse GM-CSF such that cocaine exposure increases both cytokines only in the absence of HIV infection. Together these data provide new insights into the unique neurobehavioral processes underlying HIV infection and cocaine use disorders, and further how they interact to effect immune responses.

## Introduction

An estimated 1.2 million people in the United States currently live with HIV infection (CDC, 2023). Substance use disorder (SUD) is highly co-occurring with HIV infection. Cocaine use rates among people living with HIV (PLWH) are consistently considerably higher than those observed in the general population (Mimiaga et al., 2013; Parsons et al., 2014; Azar et al., 2015; Meade et al., 2015; Hartzler et al., 2017; Liang et al., 2020; LuPone et al., 2023; NIDA, 2023). Both chronic exposure to cocaine and progressive HIV infection can independently alter behavior and immune system function and may interact to create a unique neurobehavioral niche. Cocaine use disorder is characterized by difficulty in terminating drug use, chronic relapse even after periods of abstinence and persistent drug use despite adverse consequences. A greater understanding of the immune and behavioral consequences of HIV infection can enable development of targeted therapeutics to support cessation of cocaine use.

The relationship between drug exposure and HIV infection appears to be bidirectional. For example, cocaine has been demonstrated to enhance HIV replication and promote the development of HIV associated neurocognitive disorders (HAND)(Carrico et al., 2008; Cook et al., 2008; Baum et al., 2009), to elevate levels of proinflammatory cytokines in humanized mouse models prior to HIV-1 infection, and to accelerate infection (Kim et al., 2015). In addition to cocaine effects on HIV outcomes, cocaine-related behaviors are dysregulated in HIV rodent models. A majority of the work has employed transgenic models to investigate outcomes in a cocaine conditioned place preference (CPP) paradigm which assesses the development and expression of drug-context associations. In HIV-1 Tat transgenic (Tg) mice, Tat expression potentiates cocaine CPP (Paris et al., 2014; Mediouni et al., 2015a; Zhu et al., 2022). Inducing Tat expression is also sufficient to drive the reinstatement of cocaine CPP (Paris et al., 2014; Mediouni et al., 2015) following extinction, suggesting that HIV-1 proteins can act to alter cocaine-related behavior, potentially through inflammatory effects or effects on the dopamine transporter (Miller et al., 2018). Notably, HIV-1 Tg rats show blunted cocaine preference and a failure to escalate self-administration (Bertrand et al., 2018), suggesting that there may be species or expression pattern specific differences in the effects of these proteins.

One challenge in preclinical studies of the outcomes of HIV infection is limitations in modeling progressive HIV infection. There have been advances in rodent models of HIV, including the development of the EcoHIV model (Potash et al., 2005), which enables infection of wild type mice, and humanized models that can be successfully infected with HIV-1 (Yao and Buch, 2012; Dash et al., 2021; Endsley et al., 2021; Waight et al., 2022; Baroncini et al., 2023).To investigate the effects of progressive HIV-1 infection on cocaine-related behavior and how interactions between HIV-1 and cocaine exposure impact peripheral and central inflammation, the current study utilized the CD34-NSG humanized mouse model, in which bone marrow engraftment enables humanization of the mouse immune system which can support productive HIV-1 infection. These experiments assessed cocaine-related behaviors in the CPP task, including preference, extinction, and reinstatement, and the expression of preference under conflict. The findings demonstrate that HIV-1-infected humanized mice form a preference for the cocaine-paired environment of similar magnitude to non-infected controls. However, HIV-infected mice exhibited greater preferences across extinction, indicating resistance to extinction of cocaine CPP. The results further demonstrate HIV-1 and cocaine effects on peripheral cytokine/chemokine levels, which were selective to human or mouse targets, and further, microglial and astrocyte changes in key neuroanatomical substrates of reward related memory. Together, these findings characterize HIV effects on cocaine-related behaviors and identify independent and interactive effects of HIV-1 infection and cocaine exposure on peripheral and central immune outcomes.

## Methods

### Subjects

Adult male (n = 25) and female (n = 40) NOD/Scid IL2Rg−/− (NSG) mice were used for these studies (Gorantla et al., 2010b, 2010a; Dash et al., 2011; Boska et al., 2014; Bade et al., 2016). Prior to arriving at Drexel University Animal Facilities, transgenic NOD/Scid IL2Rg−/− (NOG) mice were irradiated and underwent transplantation of human hematopoietic stem and progenitor cells (HPSCs). Blood was collected and tested prior to transportation to ensure successful engraftment. The average age upon arrival was 182 days. Following arrival, mice underwent a 2-week quarantine period, followed by placement in their final housing location under microisolation conditions at Drexel University’s Center City campus. All mice were housed in same sex cages with *ad libitum* access to food and water in a temperature and humidity-controlled environment under a standard 12-hour light/dark cycle. Mice were given supplementary diet gel as needed to maintain healthy body weights. All procedures were approved by the Institutional Animal Use and Care Committee at Drexel University.

### HIV inoculation and viral load testing

Following a 1-week acclimation, mice were matched based on sex and engraftment status into either HIV or sham control groups. Mice were inoculated with 250ng p24 of the R5-tropic HIV-1 strain ADA (HIV_ADA_). For inoculation, mice were briefly anesthetized with 5% vaporized isoflurane and injected interperitoneally with either HIV-1 or vehicle (RPMI medium) for sham controls prior to being returned to their home cage.

Blood was collected from all mice every two weeks. For blood collections, mice were briefly anesthetized with 5% vaporized isoflurane and, using the submandibular collection method, approximately 200µL of blood was collected from each mouse and spun at 9000rpm for 20 minutes at 4°C. Plasma was collected and stored at −80°C. Viral load for each mouse was determined via Q-RT-PCR performed by the Penn CFAR Viral Reservoir Core at the University of Pennsylvania. RNA was prepared using Tri reagent. Samples were run in duplicate using LTR primer-probe to determine HIV-1 RNA copies. The sequence of primers for RCAS were 5’-GTC AAT AGA GAG AGG GAT GGA CAA A-3’ and R-5’-TCC ACA AGT GTA GCA GAG CCC-3’ and the sequences for HIV-1 LTR were 5’-GCC TCA ATA AAG CTT GCC TTG A-3’ and R-5’-GGG CGC CAC TGC TAG AGA-3’. RCAS virion was spiked in each sample and amplified separately to confirm virus/RNA recovery and a lack of PCR inhibition.

### Conditioned place preference paradigm

The CPP paradigm was used to investigate cocaine reward-related behaviors. Med Associates CPP boxes were used for all experiments. These apparatuses consist of 3 distinct chambers, connected by openings which can be closed by the experimenter. The small center chamber had gray walls and a solid gray floor. On either side of the center chamber were two larger chambers, one with black walls with metal bar flooring, the other with white walls with a metal grid floor. On the first day of the CPP protocol, mice were placed in the center chamber and allowed to explore all 3 chambers freely for 20 minutes. Using Med Associates software, data collected included latency to enter each chamber, number of entries, total movement and activity, and time spent in each chamber. Data from this session were used as an initial baseline for preference. Mice were then matched based on infection status to either cocaine or saline-only control groups. Mice in the cocaine group received an i.p. injection of cocaine (10mg/kg) immediately prior to placement in the assigned reward-paired chamber. On alternating days, mice received a saline injection prior to placement in the saline-paired chamber. There was a total of 6 conditioning sessions (3 each, cocaine and saline pairings). Saline controls received saline prior to all sessions. The order of sessions was counterbalanced among groups for initial preference, drug condition, and infection status. To assess the development of a preference for the cocaine-paired chamber, mice were placed into the gray chamber and allowed to explore all 3 chambers freely for 20 (extinction and reinstatement tasks) or 5 minutes (compulsive-like cocaine seeking task).

#### Extinction and Reinstatement

In order to investigate persistent expression of a CPP after the cocaine-context relationship was removed, one cohort of mice underwent extinction training. In the 3 consecutive extinction sessions, mice were placed in the center chamber and allowed to freely explore all 3 chambers for 20 minutes in the absence of cocaine. To investigate a model of stress-primed reinstatement of cocaine seeking in HIV infection, the ability of a pharmacological stressor, yohimbine, to reinstate cocaine seeking was assessed. One day after the third extinction session, mice were administered a single dose of the α2 adrenergic antagonist yohimbine (2mg/kg) 30 minutes prior to placement in the CPP apparatus. These sessions were identical to the extinction sessions other than drug treatment.

#### Cocaine seeking under conflict

To investigate persistent cocaine-seeking despite adverse consequences, a separate cohort of mice experienced a footshock in the cocaine-paired chamber following conditioning under parameters that have been shown to suppress alcohol (Sneddon et al., 2018; Xie et al., 2019) and food (Barker et al., 2013) seeking behaviors in mice. Briefly, the CPP test consisted of a 5-minute session to avoid extinction of the learned cocaine-chamber association. 24 hours after the CPP test, mice were restricted to the cocaine-paired chamber for 3 minutes and received a mild (0.8 mA, 2 seconds) foot-shock 1 minute after placement. The following day mice underwent a 20-minute CPP test session, and the first 5 minutes were quantified to compare to the pre-shock test session.

### Brain Tissue Processing and Immunohistochemistry

Following the conclusion of all behavioral testing, mice were anesthetized, and blood was collected for viral load testing via submandibular bleeds for comparison to other time points. Mice were then overdosed on isoflurane and transcardially perfused with 1X PBS solution followed by 4% Paraformaldehyde (PFA). Brains were collected and stored in 4% PFA for 24 hours, then cryoprotected in sucrose solution for subsequent immunohistological processing for assessment of astrocyte immunoreactivity via GFAP immunohistochemistry and putative microglia number and reactivity via Iba1 immunohistochemistry.

Using a cryostat, 40µm coronal sections were collected in quadruplicate and stored in 0.01% sodium azide. Tissue sections were incubated in 1% hydrogen peroxide for 1 hour, then blocked in 5% normal donkey serum for 1 hour prior to incubation in primary antibody (anti-GFAP primary antibody, 1:10,000, G9269, Sigma-Aldrich or anti-Iba1 primary antibody 1:10,000, 019-19741, Wako) overnight at room temperature. Sections were then incubated in biotinylated donkey-anti-rabbit secondary (1:1,000, 711-065-152, Jackson labs) for 30 minutes. Staining was visualized using DAB for GFAP staining and nickel-enhanced DAB (Vector Labs, SK-4100) for Iba1 staining for 20 minutes. Sections were mounted on plus slides and coverslipped with DPX mounting medium.

For both GFAP and Iba1 analysis, 10X images were captured and stitched together using Microsoft Image Composite Editor. Images were analyzed using ImageJ software, using a thresholding approach. Number of cells (cells/area) and percent area stained were analyzed for the prefrontal cortex (PFC, 1.53mm, 1.7mm, and 1.97mm anterior of bregma), nucleus accumbens core and shell (1.54mm, 1.34mm, 1.1mm, and 0.98mm anterior of bregma), the basolateral amygdala (−0.95mm, −1.07mm, −1.23mm, and −1.31mm posterior of bregma) and dorsal hippocampus subregions (−1.3mm, −1.46mm, −2.2mm, and −2.9mm posterior of bregma) by an investigator blind to conditions.

### Cytokine and Chemokine Assessment

This study used the Luminex xMAP technology for multiplexed quantification of 48 human cytokines, chemokines, and growth factors and 32 mouse cytokines, chemokines, and growth factors. The multiplexing analysis was performed using the Luminex 200 system (Luminex, Austin, Texas) by Eve Technologies Corporation (Calgary, Alberta). Forty-eight human markers and 32 mouse markers were measured in plasma samples from week 7 or at the time of euthanasia using Eve Technologies’ Human Cytokine Panel A 48-Plex Discovery Assay and Mouse Cytokine, 32-Plex Discovery Assay (Millipore Sigma, Burlington, Massachusetts) according to the manufacturer’s protocol. Assay sensitivities of the human markers range from 0.14-50.78 pg/mL and the mouse markers range from 0.3-30.6 pg/mL. Markers included in the analysis are reported in Tables 1 and 2. A subset of targets were excluded because fewer than 5 samples per group were within the range of detection or within the standard curve.

**Table 1.** Results of the Mouse Cytokine Discovery Assay. Main effects of HIV and cocaine and interactions are represented by red, blue, and green colors, respectively.

| Analyte (units) | Abbreviation | Effect of HIV | Effect of Cocaine | Interaction |
| --- | --- | --- | --- | --- |
| Eotaxin | Eotaxin | F (1, 28) = 0.1355, p = 0.7156 | F (1, 28) = 0.7299, p = 0.4002 | F (1, 28) = 0.1563, p = 0.6956 |
| Granulocyte-colony stimulating factor | G-CSF | **F (1, 28) = 4.517, p = 0.0425** | F (1, 28) = 0.2579, p = 0.6155 | F (1, 28) = 1.193, p = 0.2840 |
| Interleukin 1 alpha | IL-1α | **F (1, 26) = 6.304, p = 0.0186** | F (1, 26) = 0.01626, p = 0.8995 | F (1, 26) = 0.01455, p = 0.9049 |
| Interleukin 6 | IL-6 | F (1, 28) = 1.420, p = 0.2433 | F (1, 28) = 2.094, p = 0.1590 | F (1, 28) = 1.340, p = 0.2568 |
| Interleukin 12 subunit | IL-12p40 | F (1, 23) = 0.8758, p = 0.3591 | F (1, 23) = 1.255, p = 0.2741 | F (1, 23) = 0.5735, p = 0.4565 |
| Interferon gamma inducible protein-10 | IP-10 | **F (1, 28) = 5.540, p = 0.0258** | F (1, 28) = 1.181, p = 0.2865 | F (1, 28) = 0.2370, p = 0.6302 |
| Interleukin 13 | IL-13 | F (1, 28) = 0.8561, p = 0.3627 | F (1, 28) = 0.5442, p = 0.4668 | F (1, 28) = 0.3758, p = 0.5448 |
| Interleukin 15 | IL-15 | F (1, 28) = 0.009636, p = 0.9225 | F (1, 28) = 0.01326, p = 0.9092 | F (1, 28) = 1.876, p = 0.1817 |
| Keratinocyte Chemoattractant, also called chemokine ligand 1 (CXCL1) | KC/CXCL1 | F (1, 28) = 0.07106, p = 0.7918 | F (1, 28) = 2.786, p = 0.1063 | F (1, 28) = 0.2033, p = 0.6555 |
| CXC Chemokine ligand 5 (CSCL5) | LIX/CSCL5 | F (1, 28) = 3.066, p = 0.0909 | F (1, 28) = 0.9750, p = 0.3319 | F (1, 28) = 0.2286, p = 0.6363 |
| Macrophage colony stimulating factor | M-CSF | F (1, 28) = 0.005562, p = 0.9411 | **F (1, 28) = 4.644, p = 0.0399** | F (1, 28) = 0.01024, p = 0.9201 |
| Monocyte chemoattractant protein-1 | MCP-1 | F (1, 28) = 1.386, p = 0.2490 | F (1, 28) = 2.019, p = 0.1664 | F (1, 28) = 0.0007655, p = 0.9781 |
| Chemokine ligand 9 (CXCL9) | MIG/CXCL9 | F (1, 28) = 2.288, p = 0.1416 | F (1, 28) = 0.1762, p = 0.6778 | F (1, 28) = 0.1359, p = 0.7152 |
| Macrophage inflammatory protein-1 alpha | MIP-1α | F (1, 28) = 0.04394, p = 0.8355 | F (1, 28) = 0.06507, p = 0.8005 | F (1, 28) = 0.05794, p = 0.8115 |
| Macrophage inflammatory protein-1 beta | MIP-1β | F (1, 28) = 0.4902, p = 0.4896 | F (1, 28) = 0.007501, p = 0.9316 | F (1, 28) = 0.4255, p = 0.5195 |
| Macrophage inflammatory protein-2 | MIP-2 | F (1, 28) = 1.365, p = 0.2525 | F (1, 28) = 0.009673, p = 0.9224 | F (1, 28) = 0.08682, p = 0.7704 |
| Regulated upon activation, normal T cell expressed and presumably secreted | RANTES/CCL5 | **F (1, 27) = 4.227, p = 0.0496** | F (1, 27) = 0.05951, p = 0.8091 | F (1, 27) = 0.7640, p = 0.3898 |
| Tumor necrosis factor alpha | TNFα | F (1, 28) = 1.250, p = 0.2730 | F (1, 28) = 0.2502, p = 0.6208 | F (1, 28) = 0.02377, p = 0.8786 |

### Statistical Analysis

All data were analyzed in GraphPad. Behavioral data were analyzed by repeated measures ANOVA for 2-way comparisons or by unpaired t-tests. Cytokine/chemokine expression data and immunohistological data were analyzed with 2way ANOVA. Analyses were Greenhouse-Geisser corrected when sphericity was not met. Significant interactions were followed with post hoc analyses.

## Results

### HIV-1 infection of humanized mice

Adult male and female NOD/Scid IL2Rg−/− (NSG) mice were matched by engraftment status to undergo either sham or HIV inoculation and cocaine/saline exposure. There were no differences in initial human monocyte or lymphocyte composition between groups of mice, as confirmed by assessing expression of CD3 [HIV groups: F (1,61) = 2.216, p = 0.1417; cocaine exposure: F (1,61) = 1.381, p = 0.2444; **Fig 1A**], CD14 [HIV groups: F (1,61) = 0.7720; p = 0.3831; cocaine exposure: F (1,61) = 2.558, p = 0.1149], and CD19 [HIV groups: F (1,61) = 1.910, p = 0.1720; cocaine exposure: F (1,61) = 1.346, p = 0.2505].

**Figure 1.**
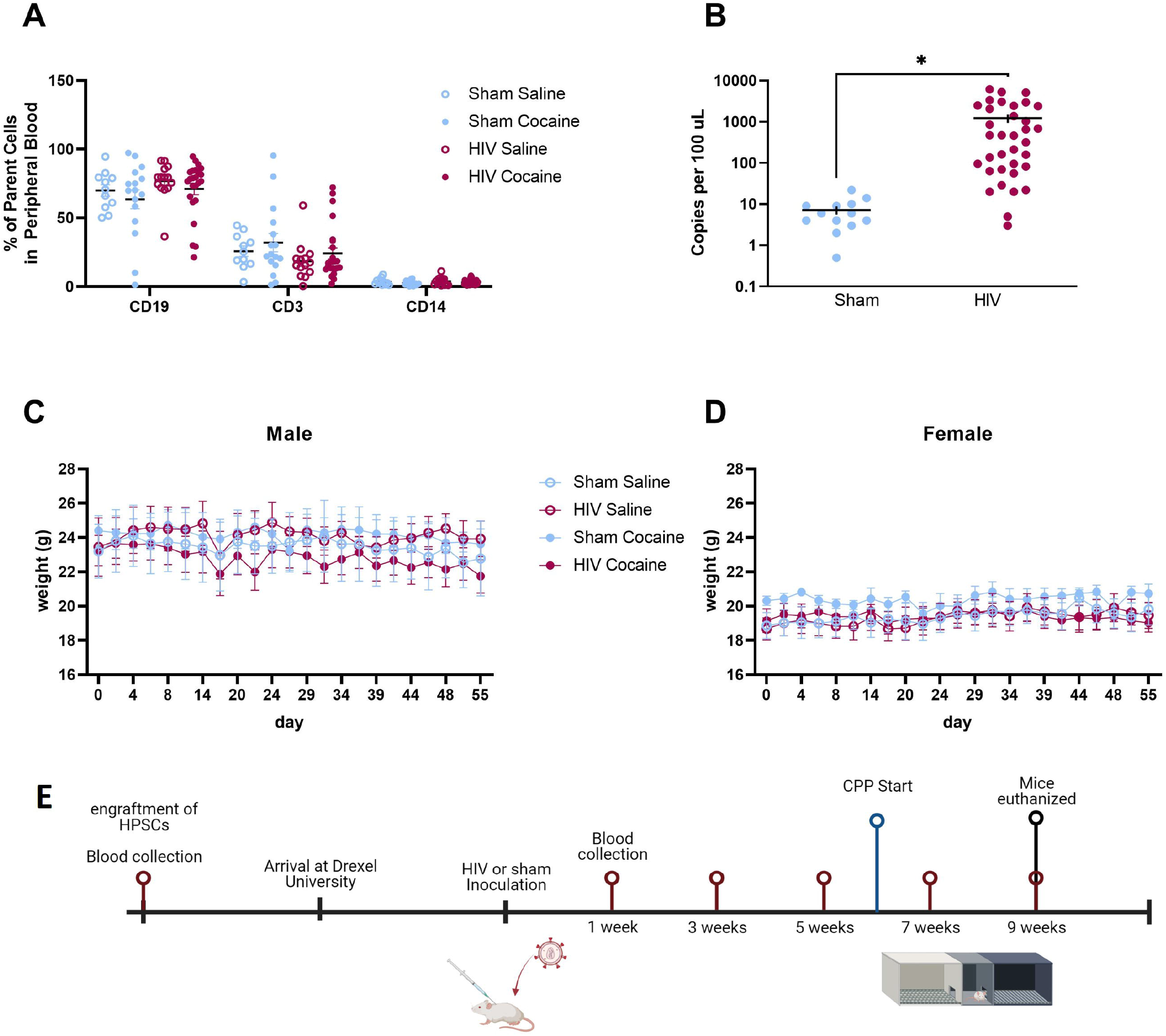
Humanized mouse model of HIV-1 infection. (A) Mice were matched based on engraftment status to undergo HIV-1 vs sham infection and subsequent cocaine vs saline treatment. (B) The number of copies of HIV-1 RNA per 100µl of plasma sample was elevated in HIV-1 inoculated mice (n = 26) compared to Sham (n = 11) mice. (C) No effect of HIV-1 infection or cocaine treatment were observed on weight (grams) of male mice (n = 24) or female (D) mice throughout the duration of the experiment. (D) (E) Timeline of experiments (created with Biorender.com). *p < 0.05. Points represent mean +/− SEM.

To determine HIV viral load, HIV-1 RNA was measured using Q-RT-PCR. An unpaired t-test of RNA copies was performed to compare viral load in HIV and sham inoculated mice. There was a significant difference in HIV viral load between HIV inoculated mice and sham inoculated mice [t(46) = 2.591, p = 0.0128; **Fig 1B**].

Mouse weights were collected three times per week across the study. In male cohorts, three-way ANOVA demonstrated no effects of HIV [F(1,20) = 0.09260, p = 0.7640], cocaine [F(1,20) = 0.06748, p = 0.7977], or time [F(5.494,109.9) = 2.227, p = 0.0509; **Fig 1C**]. There were no effects of HIV [F(1,27) = 0.07297, p = 0.4005] or cocaine [F(1,27) = 0.6618, p = 0.4230] on female mouse weights, however there was a main effect of time [three-way ANOVA; F(9.966,188.1) = 2.783, p = 0.0090; Greenhouse-Geisser corrected; **Fig 1D**]. Post hoc comparisons revealed that weights were greater than day 0 on days 2, 29, 32, 36, 39, and 47 (p’s < 0.05).

### HIV infection does not alter cocaine preference in mice

In order to assess preference for the cocaine-paired chamber, mice were trained in alternating sessions to associate distinct environments with cocaine or saline. Across training, mice exhibited higher locomotion during cocaine sessions than saline sessions [two-way ANOVA F(1,25) = 94.23, p < 0.0001; **Fig 2A**]. There was no effect of HIV infection on locomotion [two-way ANOVA; F(1,25) = 2.590, p = 0.1201] and no interaction [F(1,25) = 0.006569, p = 0.9360]. Following training, mice were tested for a cocaine preference in a 20-minute session in which they were allowed to freely explore the CPP apparatus. CPP score was determined by comparing time spent in the cocaine-paired and unpaired chambers (time in paired – unpaired). Compared to baseline scores, all mice demonstrated a preference for the cocaine-paired chamber [two-way ANOVA F(1,25) = 20.82, p = 0.0001; **Fig 2B**]. HIV infection did not impact the formation of a preference for the cocaine-paired chamber [two-way ANOVA; F(1,25) = 0.06663, p = 0.4220]. Together these data demonstrate that HIV infection did not impact acquisition or expression of cocaine conditioned place preference.

**Figure 2.**
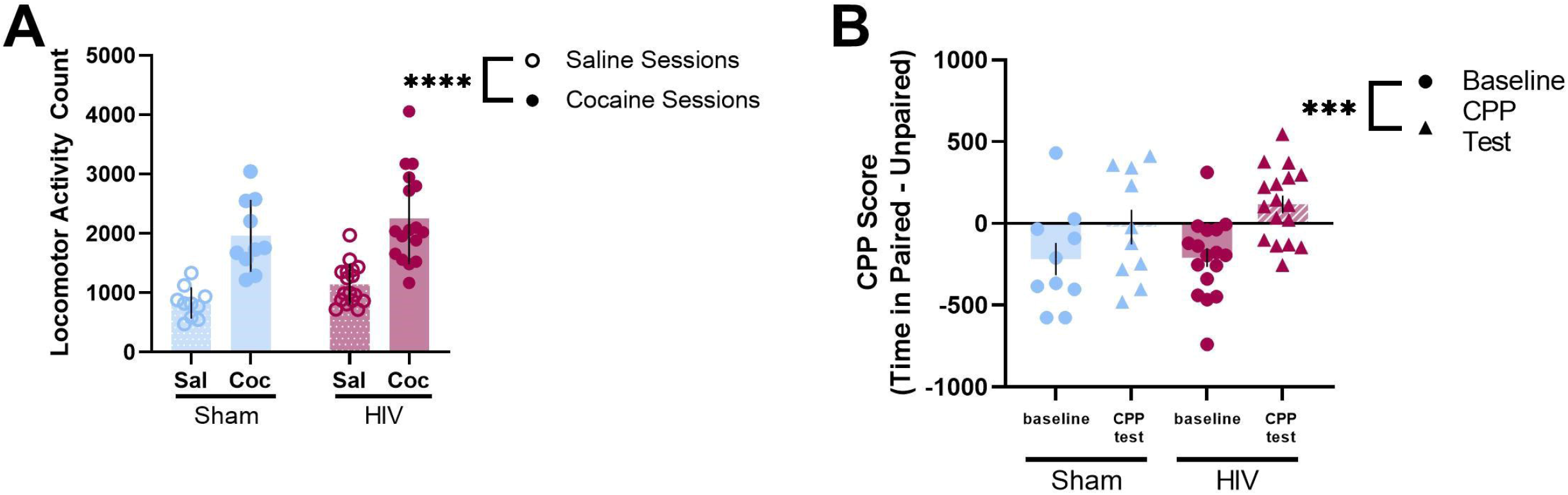
Normal cocaine-induced locomotion and conditioned place preference in HIV-1 infected mice. (A) Average locomotor scores of cocaine-exposed sham (n=10) or HIV-infected (n=17) mice during saline or cocaine-paired conditioning sessions. (B) No differences in CPP scores were observed in cocaine-exposed sham or HIV-infected mice. ***p < 0.001, ****p<0.0001. Bars represent mean +/− SEM.

### HIV infection promotes persistent cocaine seeking across extinction and reinstatement

To determine the effect of HIV infection on extinction and stress-induced reinstatement of cocaine seeking mice were allowed three sessions without cocaine followed by a session in which mice were administered yohimbine prior to testing (**Fig 3A**). HIV infection inhibited extinction such that HIV infected mice continued to spend more time in the cocaine-paired chamber than sham-infected counter parts [two-way ANOVA; F(1,24) = 4.913, p = 0.0364 **Fig 3B**]. A main effect of time was observed across sessions [F(2.723, 65.34) = 5.225, p = 0.0036; Greenhouse-Geisser corrected].

**Figure 3.**
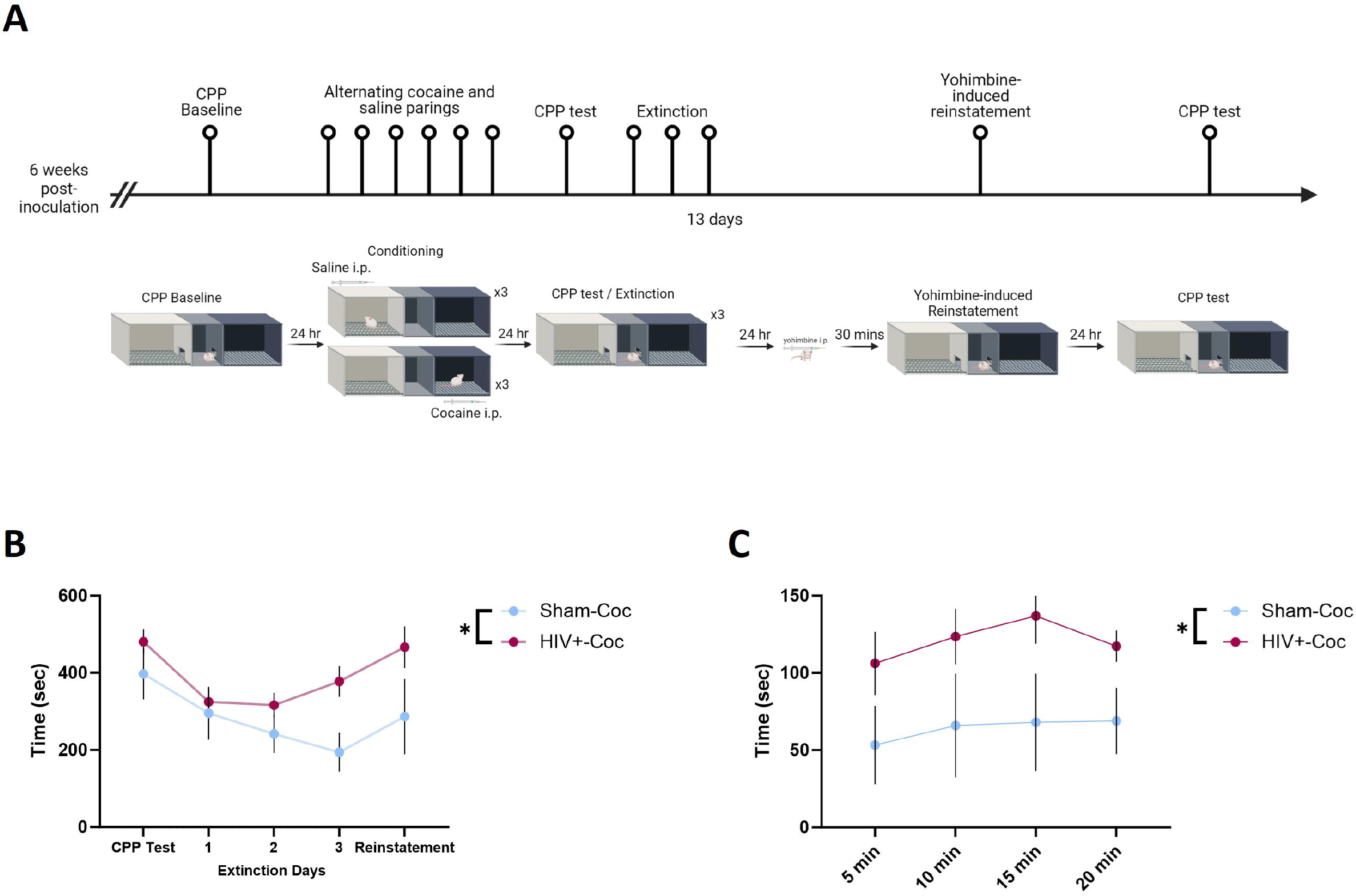
Impaired extinction of cocaine CPP in HIV-1 infected mice. (A) Timeline of CPP followed by extinction and reinstatement testing (created with Biorender.com). (B) A subset of mice underwent extinction of cocaine CPP following conditioning. HIV-1 infected mice exhibited persistent CPP across extinction sessions, indicative of a deficit in extinction. (C) Sham- and HIV-1 infected mice exhibited similar magnitudes of yohimbine-induced reinstatement. *p < 0.05. Data represent mean +/− SEM.

Time spent in the cocaine-paired chamber during the reinstatement test differed significantly between sham and HIV infected mice [two-way ANOVA; F (1,23) = 4.789, p = 0.0391; **Fig 3C**]. A main effect of time was also observed [F (2.149, 49.42) = 0.9069, p = 0.0416; Greenhouse-Geisser corrected], but no interaction between HIV and time was observed [F (3, 69) = 0.1961, p = 0.8987].

### Sham and HIV-infected mice reduce cocaine seeking under conflict

To assess whether HIV infection impacted cocaine seeking despite adverse consequences, time spent in the reward paired chamber before and after a footshock were compared (**Fig 4A**). A main effect of time was observed such that all cocaine-exposed mice reduced time in the cocaine-paired chamber following footshock [two-way ANOVA; F (1,9) = 16.49, p = 0.0028; **Fig 4B**]. There was no effect of HIV [F (1,9) = 1.945, p = 0.1966] and no interaction [F (1,9) = 0.4919, p = 0.5008]. Latency to enter the cocaine-paired chamber was not altered after footshock [two-way ANOVA; F (1,9) = 3.082, p = 0.1130]. There was no effect of HIV [F (1,9) = 0.3797, p = 0.5530] and no interaction [F (1,9) = 0.2538, p = 0.6262] observed (**Fig 4C**).

**Figure 4.**
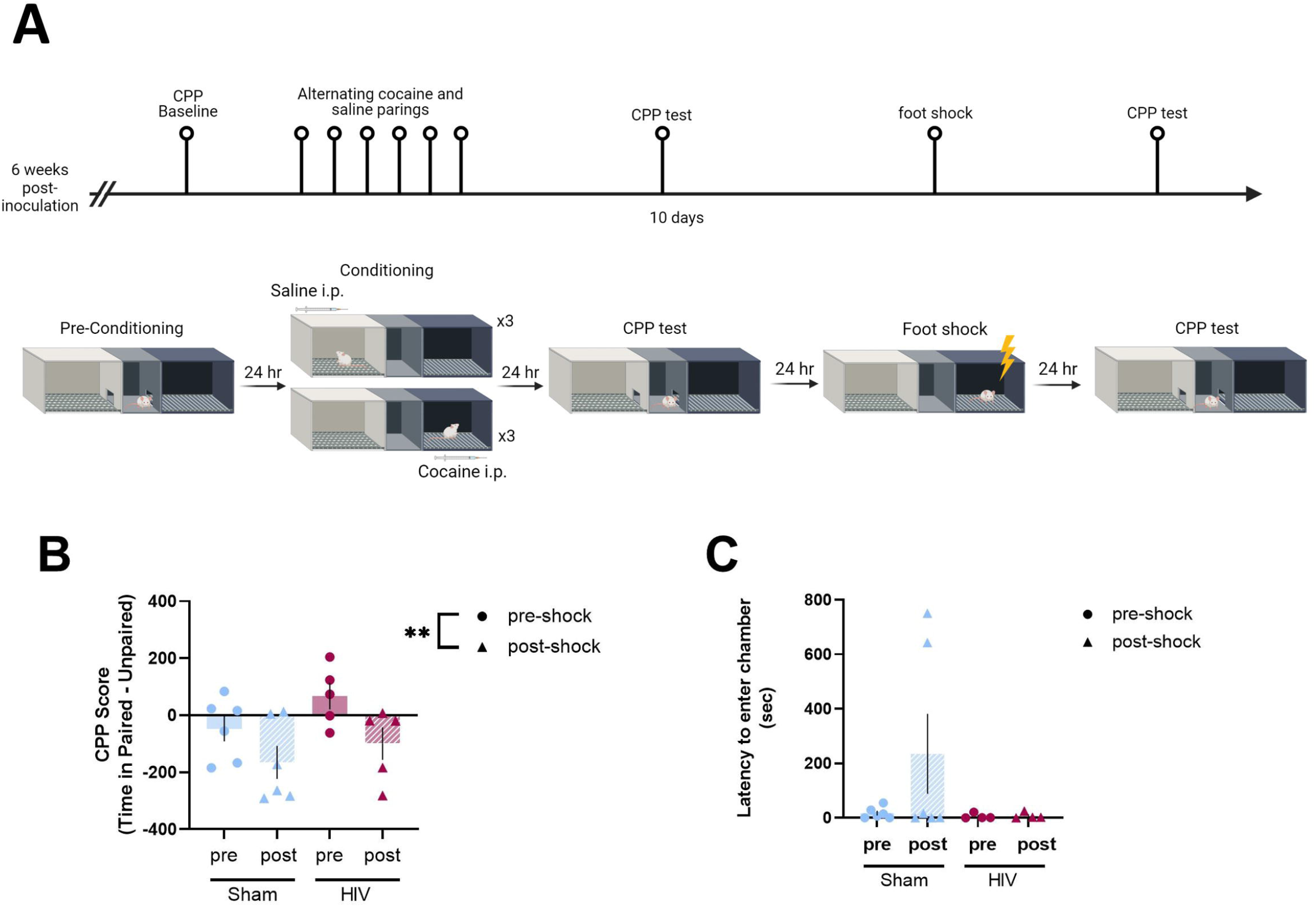
HIV infection did not impact cocaine seeking under conflict. (A) Timeline of CPP with footshock and testing for cocaine seeking under conflict. (B) Sham (n=6) and HIV-infected mice (n=5) showed similar reductions of time spent in the cocaine-paired chamber when cocaine-footshock conflict was introduced. (B) Neither sham (n=6) and HIV-infected mice (n=5) showed significant increases in latencies to enter the cocaine-paired chamber following the introduction of cocaine-footshock conflict. **p < 0.01. Data represent mean +/− SEM.

### GFAP immunoreactivity is increased in the PFC of HIV-infected mice

To determine if HIV infection or cocaine exposure altered GFAP immunoreactivity, brain tissue was analyzed for the percent area of GFAP staining in reward substrates (**Fig 5A**). A two-way ANOVA revealed an increased percent area of GFAP staining in HIV-infected mice in the prelimbic [main effect of HIV: F (1,29) = 5.619, p = 0.0249; **Fig 5B**] and infralimbic [main effect of HIV: F (1,28) = 4.447, p = 0.0440; **Fig 5C**] PFC, but no effect of cocaine [PrL: F (1,28) = 1.733, p = 0.1998, IL: F (1,28) = 0.568, p = 0.4572] and no interaction [PrL: F (1,28) 0.3613, p = 0.5526, IL: F (1, 28) = 0.374, p = 0.5455] were observed.

**Figure 5.**
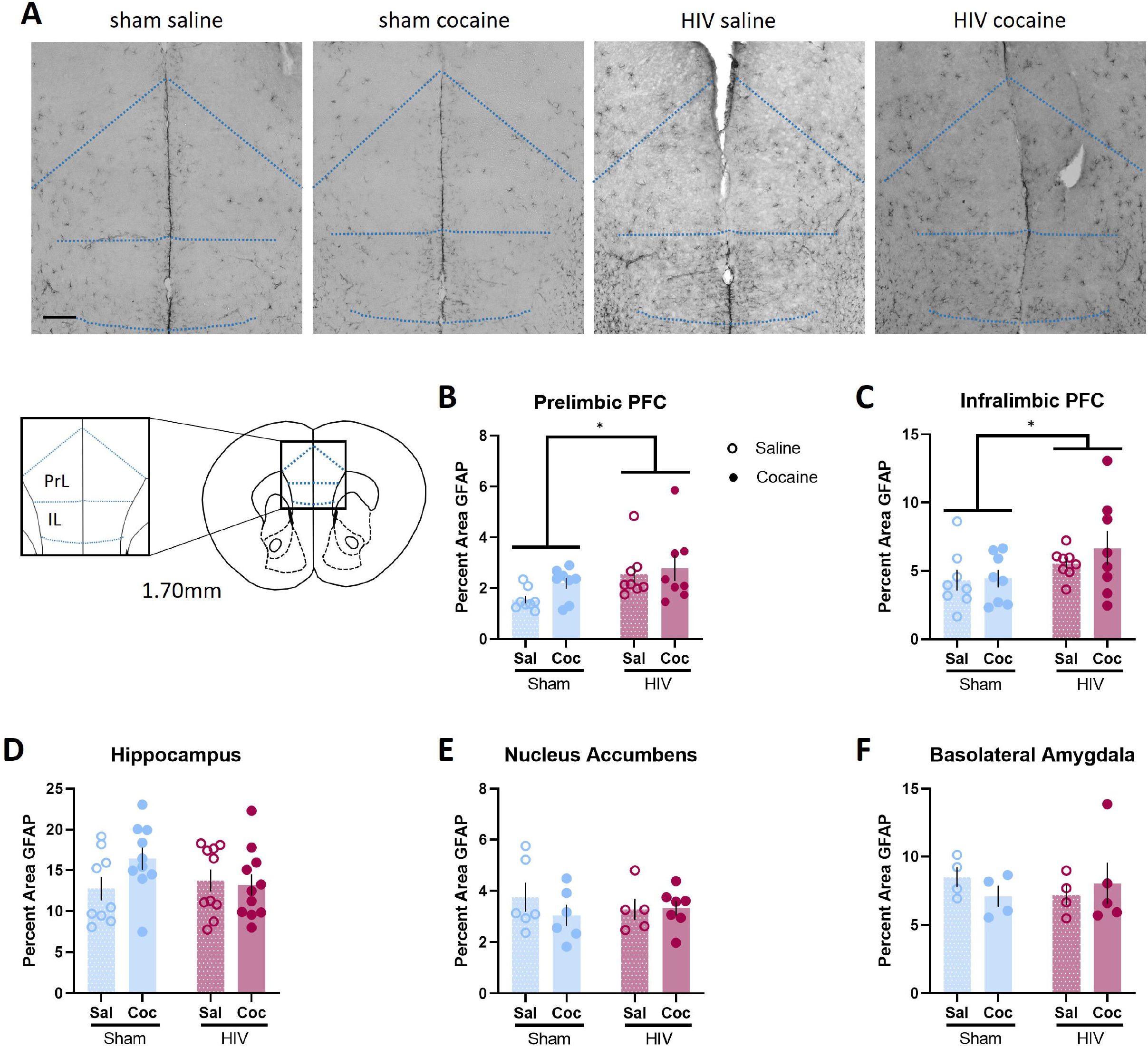
HIV-1 infection altered GFAP immunoreactivity in reward-related neural substrates. (A) Representative images of GFAP staining in the PFC from sham-saline, sham-cocaine, HIV+-saline, and HIV+-cocaine mice. (B, C) HIV-1 infected mice exhibited increased GFAP immunoreactivity in the infralimbic and prelimbic subregions of the PFC. (D, E, F). No effects of HIV-1 infection or cocaine exposure on GFAP immunoreactivity in the (D) hippocampus, (E) nucleus accumbens, or (F) basolateral amygdala were observed. *p < 0.05. Data represent mean +/− SEM. Scale bar represents 250µm.

No effects of cocaine or HIV infection on GFAP immunoreactivity were observed in the hippocampus [interaction: F (1,36) = 2.382, p = 0.1315; main effect of cocaine: F (1,36) = 1.322, p = 0.2578; main effect of HIV: F (1,36) = 0.6809, p = 0.4147; **Fig 5D**], the nucleus accumbens [interaction: F (1,20) = 0.7723, p = 0.3900; main effect of cocaine: F (1,20) = 0.5910, p = 0.4510; main effect of HIV: F (1,20) = 0.04610, p = .08322; **Fig 5E**], or the basolateral amygdala [interaction: F (1,13) = 1.011, p = 0.3330; main effect of cocaine: F (1,13) = 0.06364, p = 0.8048; main effect of HIV: F (1,13) = 0.0269, p = 0.8722; **Fig 5F**].

### Cocaine exposure increased microglia number in the hippocampus of HIV-infected mice

To assess whether HIV infection or cocaine exposure altered the number of microglia, brain tissue was stained for detection of Iba1 and the number of Iba1+ cells was analyzed.

The hippocampus was analyzed by the four distinct subregions, the CA1 [**Fig 6A**], CA2, CA3, and dentate gyrus (DG). A two-way ANOVA revealed a significant interaction of cocaine exposure and HIV infection on the number of Iba1 cells in the CA1 subregion of the hippocampus [F (1,27) = 4.702, p = 0.0391; **Fig 6A, 6B**]. Tukey’s HSD test for multiple comparisons found that the mean number of Iba1+ cells was significantly increased in cocaine-exposed HIV-infected mice, compared to saline-only HIV-infected mice [p = 0.0214, 95% C.I. −566.2 to –36.24].

**Figure 6.**
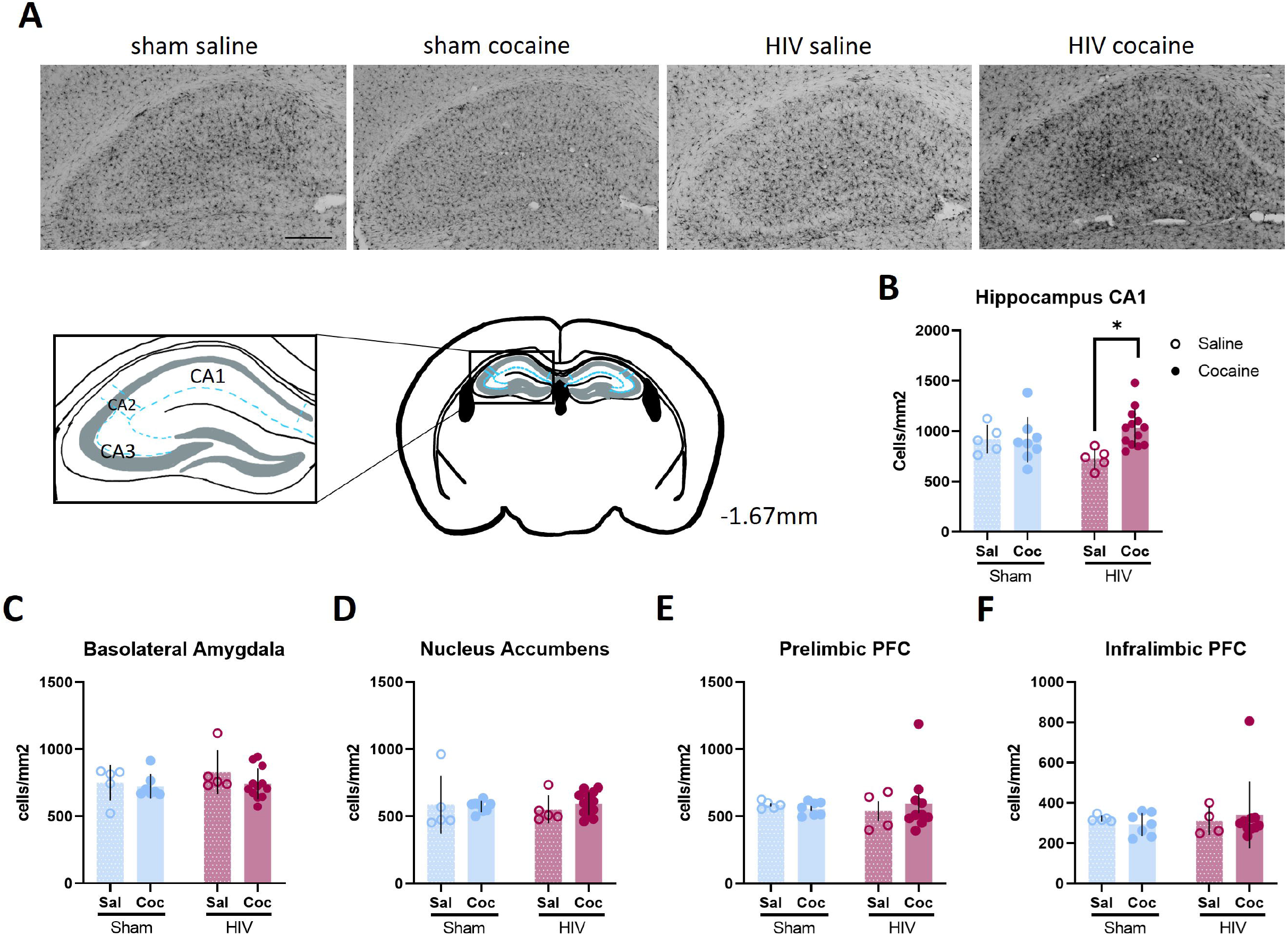
HIV infection and cocaine exposure interacted to impact microglial number in the hippocampus. (A) Representative 20X images of IBA1 staining in the hippocampus of sham-saline, sham-cocaine, HIV-saline, and HIV-cocaine mice. (B) A greater number of IBA1+ cells per mm^2^ were observed in the CA1 subfield of the hippocampus in HIV-1 infected mice exposed to cocaine. No effect of cocaine was observed in sham mice. (C, D, E, F) No effects of HIV-1 infection or cocaine exposure on microglia numbers were observed in the (C) basolateral amygdala, (D) NAc, (E) prelimbic PFC, or (F) infralimbic PFC. *p < 0.05. Data represent mean +/− SEM. Scale bar represents 250µm.

Iba1+ cell counts were not significantly different in the CA2, CA3, or DG subregions of the hippocampus (**Supplemental Fig 1A-C**). Similarly, no changes in Iba1+ cell counts were observed in the basolateral amygdala [interaction: F (1,25) = 0.4168, p = 0.5244; main effect of cocaine: F (1,25) = 1.317, p = 0.2620; main effect of HIV: F (1,25) = 0.9903, p = 0.3292; **Fig 6C**], the nucleus accumbens [interaction: F (1,27) = 0.4037, p = 0.5306; main effect of cocaine: F (1,27) = 0.1133, p = 0.7390; main effect of HIV: F (1,27) = 0.02305, p = 0.8805; **Fig 6D**], the prelimbic PFC [interaction: F (1,22) = 0.4181, p = 0.5246; main effect of cocaine: F (1,22) = 0.0473, p – 0.8297; main effect of HIV: F (1,22) = 0.0051, p = 0.9436; **Fig 6E**], or the infralimbic PFC [interaction: F (1,22) = 0.3847, p = 0.5415; main effect of cocaine: F (1,22) = 0.0002, p = 0.9872; main effect of HIV: F (1,22) = 0.1507, p = 0.7016; **Fig 6F**].

### Microglial activation is elevated in cocaine-exposed HIV-infected mice

To determine if microglial activation is changed by cocaine exposure or HIV infection, the percent area of Iba1+ cells was analyzed (**Fig 7A**). A two-way ANOVA showed a main effect of cocaine on the percent area of Iba1+ cells in the basolateral amygdala [F (1, 25) = 4.468, p = 0.0447; **Fig 7B**]. No effect of HIV [F (1,25) = 0.5165, p = 0.4790] or interaction [F (1,25) = 0.5651, p = 0.4592] was observed.

**Figure 7.**
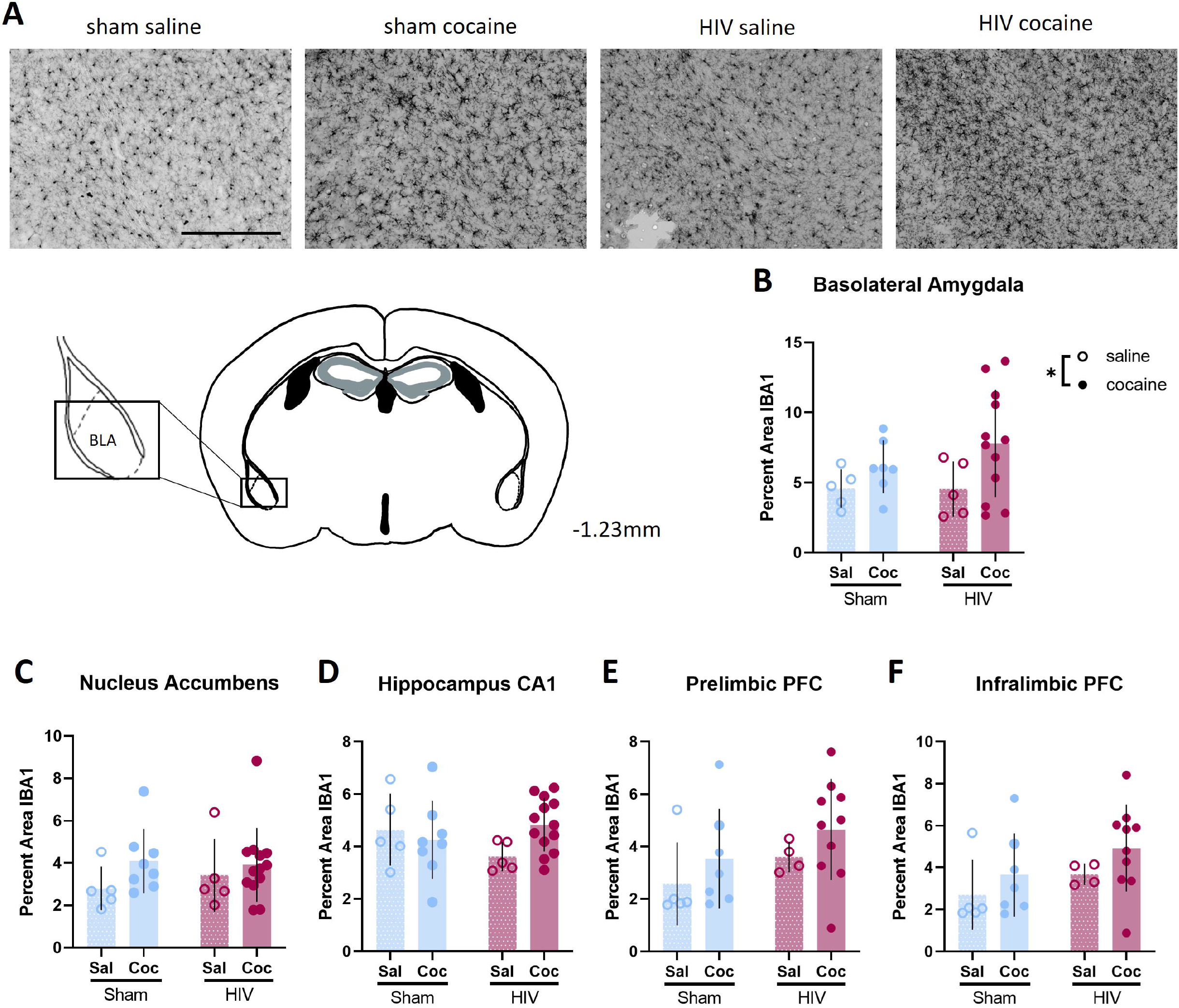
Cocaine altered microglial activation in the basolateral amygdala. (A) Representative 20X images of IBA-1 staining in the BLA (X bregma) from sham-saline, sham-cocaine, HIV-saline, and HIV-cocaine mice. (B) Independent of HIV-1 infection status, cocaine exposure increased the percent area of IBA1 staining in the BLA. (C, D, E, F) No effects of HIV-1 infection or cocaine exposure were observed in the (C) NAc, (D) CA1 region of the hippocampus, (E) prelimbic PFC, or (F) infralimbic PFC. *p < 0.05. Data represent mean +/− SEM. Scale bar represents 250µm.

No changes in percent area of Iba1+ cells were observed in the CA1 [Interaction: F (1,27) = 3.152, p = 0.0871; main effect of cocaine: F (1,27) = 0.8341, p = 0.3692; main effect –of HIV: F (1,27) = 0.2408, p = 0.6276; **Fig 7C**], CA2, CA3, or DG subregions of the hippocampus (**Supplemental Fig 1D-F**) IBA1+ percent area was unchanged in the nucleus accumbens [interaction: F (1,27) = 0.4174, p = 0.5238; main effect of cocaine: F (1,27) = 2.074, p = 0.1613; main effect of HIV: F (1,27) = 0.1278, p = 0.7235; **Fig 7B**], the prelimbic PFC [interaction: F (1,22) = 0.0043, p = 0.9479; main effect of cocaine: F (1,22) = 1.944, p = 0.1772; main effect of HIV: F (1,22) = 2.170, p = 0.1549; **Fig 7E**], and infralimbic PFC [interaction: F (1,22) = 0.0391, p = 0.8450; main effect of cocaine: F (1,22) = 2.093, p = 0.1621; main effect of HIV: F (1,22) = 2.171, p = 0.1548; **Fig 7F**].

### Cytokine/Chemokine Discovery Assays

To assess markers of inflammation in the humanized HIV-1 model, this study utilized mouse (**Table 1**) and human (**Table 2**) cytokine discovery assays.

**Table 2.** Results of the Human Cytokine Discovery Assay. Main effects of HIV and cocaine and interactions are represented by red, blue, and green colors, respectively.

| Analyte (units) | Abbreviation | Effect of HIV | Effect of Cocaine | Interaction |
| --- | --- | --- | --- | --- |
| Fibroblast growth factor 2 | FGF-2 | F (1, 33) = 0.4190, p = 0.5219 | F (1, 33) = 0.09165, p = 0.7540 | F (1, 33) = 0.9378, p = 0.3399 |
| Granulocyte-macrophage colony-stimulating factor | GM-CSF | **F (1, 34) = 18.48, p = 0.0001** | **F (1, 34) = 13.22, p = 0.0009** | **F (1, 34) = 14.32, p = 0.0006** |
| Human interferon alpha-2 | IFNα2 | **F (1, 34) = 9.291, p = 0.0044** | F (1, 34) = 0.7824, p = 0.3826 | F (1, 34) = 0.8127, p = 0.3737 |
| Interferon gamma | IFNγ | **F (1, 33) = 16.91, p = 0.0002** | **F (1, 33) = 12.11, p = 0.0014** | **F (1, 33) = 10.30, p = 0.0030** |
| Interferon gamma inducible protein-10 | IP-10 | F (1, 34) = 3.976, p = 0.0542 | F (1, 34) = 1.532, p = 0.2243 | F (1, 34) = 2.456, p = 0.1263 |
| Interleukin 1-alpha | IL-1α | F (1, 34) = 0.4859, p = 0.4905 | F (1, 34) = 0.6093, p = 0.4404 | F (1, 34) = 0.9878, p = 0.3273 |
| Interleukin 2 | IL-2 | F (1, 27) = 2.122, p = 0.1567 | F (1, 27) = 0.03377, p = 0.8556 | F (1, 27) = 0.2524, p = 0.6194 |
| Interleukin 3 | IL-3 | F (1, 33) = 0.3799, p = 0.5419 | F (1, 33) = 0.8427, p = 0.3653 | F (1, 33) = 1.265, p = 0.2688 |
| Interleukin 4 | IL-4 | F (1, 33) = 0.1552, p = 0.6962 | F (1, 33) = 0.1902, p = 0.6656 | F (1, 33) = 0.6155, p = 0.4383 |
| Interleukin 5 | IL-5 | F (1, 33) = 3.226, p = 0.0816 | F (1, 33) = 2.830, p = 0.1019 | F (1, 33) = 0.2105, p = 0.6494 |
| Interleukin 6 | IL-6 | F (1, 34) = 3.100, p = 0.0873 | F (1, 34) = 1.009, p = 0.3223 | F (1, 34) = 1.013, p = 0.3214 |
| Interleukin 7 | IL-7 | F (1, 34) = 0.2483, p = 0.3215 | F (1, 34) = 0.8068, p = 0.3754 | F (1, 34) = 1.517, p = 0.2265 |
| Interleukin 12 subunit | IL-12p40 | **F (1, 30) = 11.82, p = 0.0017** | F (1, 30) = 0.8741, p = 0.3573 | F (1, 30) = 7.646e-005, p = 0.9931 |
| Interleukin 13 | IL-13 | F (1, 29) = 2.525, p = 0.1229 | F (1, 29) = 0.0006673, p = 0.9796 | F (1, 29) = 3.631, p = 0.0667 |
| Interleukin 22 | IL-22 | F (1, 33) = 1.074, p = 0.3076 | F (1, 33) = 1.595, p = 0.2155 | F (1, 33) = 0.3613, p = 0.5519 |
| Interleukin 25 | IL-17E/IL-25 | F (1, 31) = 0.2435, p = 0.6252 | F (1, 31) = 0.6598, p = 0.4228 | F (1, 31) = 0.6996, p = 0.4093 |
| Interleukin 27 | IL-27 | F (1, 29) = 0.005823, p = 0.9397 | F (1, 29) = 0.08394, p = 0.7741 | F (1, 29) = 0.6026, p = 0.4439 |
| Macrophage inflammatory protein-1 alpha | MIP-1α | F (1, 34) = 0.9433, p = 0.3383 | F (1, 34) = 0.4064, p = 0.5281 | F (1, 34) = 1.342, p = 0.2548 |
| Macrophage inflammatory protein-1 beta | MIP-1β | **F (1, 34) = 5.964, p = 0.0200** | F (1, 34) = 1.928, p = 0.1740 | F (1, 34) = 1.779, p = 0.1911 |
| Macrophage-derived chemokine | MDC | **F (1, 34) = 13.14, p = 0.0009** | F (1, 34) = 0.1361, p = 0.7145 | F (1, 34) = 0.03766, p = 0.8473 |
| Monocyte Chemoattractant Protein-1, now called chemokine ligand 2 (CCL2) | MCP-1 | F (1, 28) = 2.764, p = 0.1076 | F (1, 28) = 1.165, p = 0.2896 | F (1, 28) = 1.146, p = 0.2935 |
| Monocyte Chemoattractant Protein-3, now called chemokine ligand 7 (CCL7) | MCP-3 | F (1, 33) = 3.224, p = 0.0818 | F (1, 33) = 0.9381, p = 0.3398 | F (1, 33) = 1.294, p = 0.2635 |
| Monokine induced by gamma interferon, Chemokine ligand 9 | MIG/CXCL9 | **F (1, 31) = 11.96, p = 0.0016** | F (1, 31) = 1.221, p = 0.2778 | F (1, 31) = 0.7837, p = 0.3828 |
| Platelet-derived growth factor AA | PDGF-AA | **F (1, 30) = 8.870, p = 0.0057** | F (1, 30) = 1.309, p = 0.2616 | F (1, 30) = 2.958, p = 0.0958 |
| Platelet-derived growth factor AB/BB | PDGF-AB/BB | F (1, 34) = 4.111, p = 0.0505 | F (1, 34) = 1.338, p = 0.2554 | F (1, 34) = 0.01232, p = 0.9123 |
| Tumor necrosis factor alpha | TNFα | **F (1, 34) = 11.33, p = 0.0019** | **F (1, 34) = 4.289, p = 0.0460** | F (1, 34) = 3.492, p = 0.0703 |

Two-way ANOVA revealed changes in mouse inflammatory markers in HIV-1 infected mice. HIV-1 infected-mice exhibited reduced IL-1α [main effect of HIV: F (1,26) = 6.304, p = 0.0186; main effect of cocaine: F (1,26) = 0.0162, p = 0.8995; interaction: F (1,26) = 0.0145, p = 0.9049; **Fig 8A**] and G-CSF [main effect of HIV: F (1, 28) = 4.517, p = 0.0425; main effect of cocaine: F (1,28) = 0.2579, p = 0.6155; interaction: F (1,28) = 1.193, p = 0.2840; **Fig 8B**). In contrast, HIV-1 infected mice had higher levels of RANTES/CCL5 [main effect of HIV: F (1,27) = 4.227, p = 0.0496; main effect of cocaine: F (1,27) = 0.0595, p = 0.8091; interaction: F (1,27) = 0.7640, p = 0.3898; **Fig 8C**] and IP-10/CXCL10 [main effect of HIV: F (1,28) = 5.540, p = 0.0258; main effect of cocaine: F (1,28) = 1.181, p = 0.2865; interaction: F (1,28) = 0.2370, p = 0.6302; **Fig 8D**] compared to controls. M-CSF was not impacted by HIV-1 infection [F (1,28) = 0.0055, p = 0.9411], but was reduced in cocaine-exposed mice [F (1, 28) = 4.644, p = 0.0399; **Fig 8E**]. No HIV x cocaine interaction [F (1,28) = 0.0102, p = 0.9201] was observed. In the human cytokine assay, two-way ANOVA revealed changes in inflammatory markers. HIV-1 infection reduced expression of IL-12p40 [main effect of HIV: F (1,30) = 11.82, p = 0.0017; main effect of cocaine: F (1,30) = 0.8741, p = 0.3575; interaction: F (1,30) = 7.646, p = 0.9931; **Fig 9A**], MIG/CXCL9 [main effect of HIV: F (1,31) = 11.96, p = 0.0016; main effect of cocaine: F (1,31) = 1.221, p = 0.2778; interaction: F (1,31) = 0.7837, p = 0.3828; **Fig 9B**], MDC [main effect of HIV: F (1,34) = 13.14, p = 0.0009; main effect of cocaine: F (1,34) = 0.1361, p = 0.7145; interaction: F (1,34) = 0.0376; **Fig 9C**] and MIP-1β [main effect of HIV: F (1,34) = 5.964, p = 0.0200; main effect of cocaine: F (1,34) = 1.928, p = 0.1740; interaction: F (1,34) = 1.779, p = 0.1911; **Fig 9D**] and PDGF-AA [main effect of HIV: F (1,30) = 8.870, p = 0.0057; main effect of cocaine: F (1,30) = 1.309, p = 0.2616; interaction: F (1,30) = 2.958, p = 0.0958; **Fig 9E**]. INFα2 levels were increased in HIV-1 infected mice compared to controls [main effect of HIV: F (1,34) = 9.291, p = 0.0044; main effect of cocaine: F (1,34) = 0.7824, p = 0.3826; interaction: F (1,34) = 0.8127, p = 0.3737; **Fig 9F**]. Analysis of human TNFα revealed main effects of HIV [F (1,34) = 4.289, p = 0.0460] and cocaine [F (1,34) = 4.289, p = 0.0460], but no interaction [F (1,34) = 3.492, p = 0.0703; **Fig 9G**]. Interactions were observed between HIV infection and cocaine exposure in human IFNɣ and GM-CSF. IFNɣ was increased in sham mice exposed to cocaine compared to control sham mice [p = 0.003], HIV infected control mice and cocaine-exposed mice (p’s < 0.0001; **Fig 9H**). GM-CSF was increased in cocaine-exposed sham mice compared to all other groups (p’s <0.0001; **Fig 9I**). There were not sufficient samples with detectable levels of RANTES/CCL5 across all groups for statistical comparison of means. However, the frequency of presence of detectable levels of RANTES/CCL5 was significantly higher in HIV-1 infected mouse samples than sham mouse samples [X^2^ (1, N = 38) = 6.6037, p = 0.0101], consistent with potentially elevated levels of RANTES/CCL5 in the HIV-1 group compared to the sham mice.

**Figure 8.**
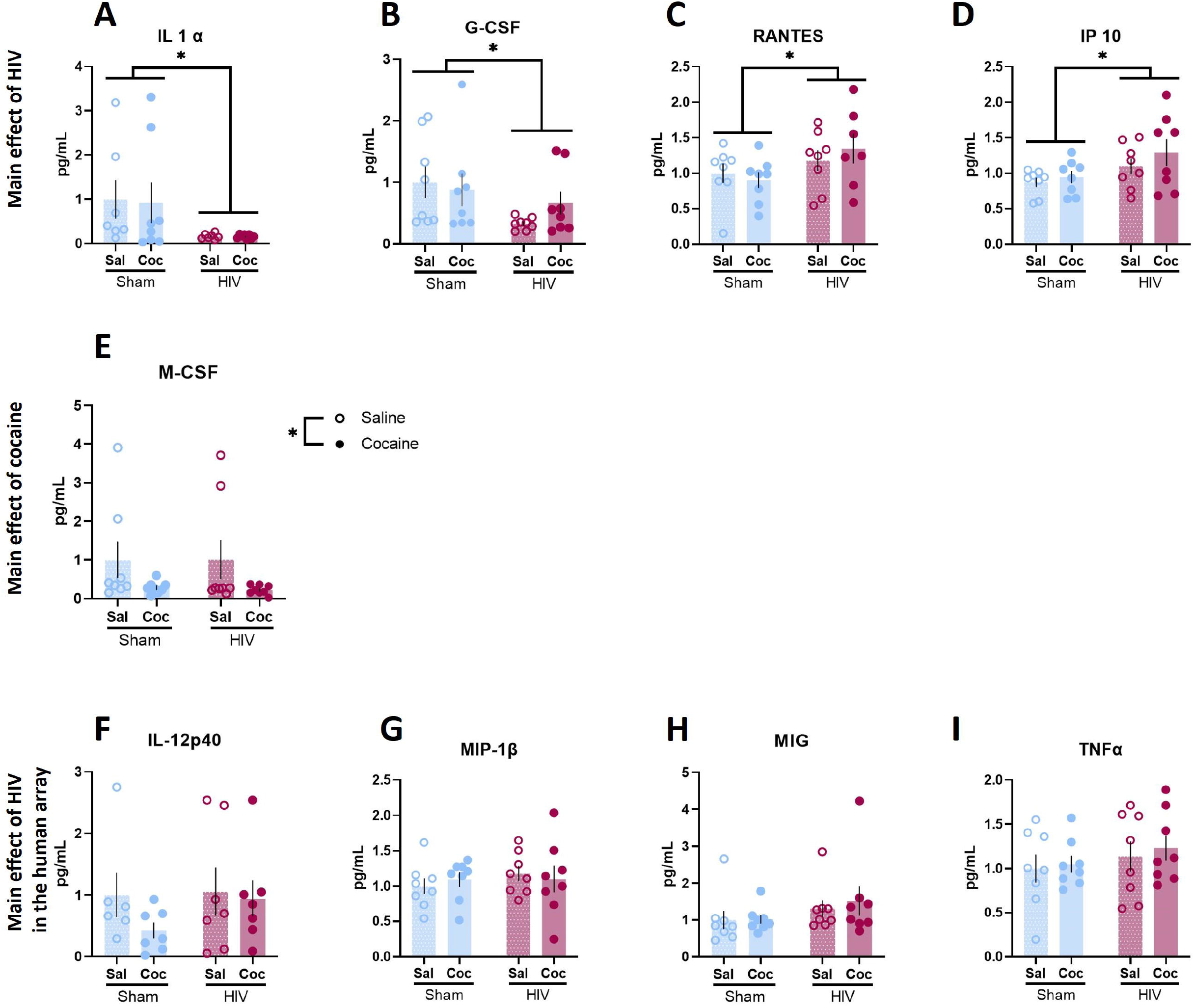
HIV and cocaine impacted mouse inflammatory factors in peripheral blood. (A, B, C, D) HIV-1 infected mice expressed lower (A) IL1α, (B) G-CSF in the periphery and increased (C) RANTES, and (D) IP 10. (E) Lower M-CSF was observed in the peripheral blood of cocaine-exposed mice. (F, G, H, I) A subset of human targets was significantly altered by HIV-1 or cocaine, while no effects were observed in the mouse inflammatory factors. *p < 0.05. Data represent mean +/− SEM.

**Figure 9.**
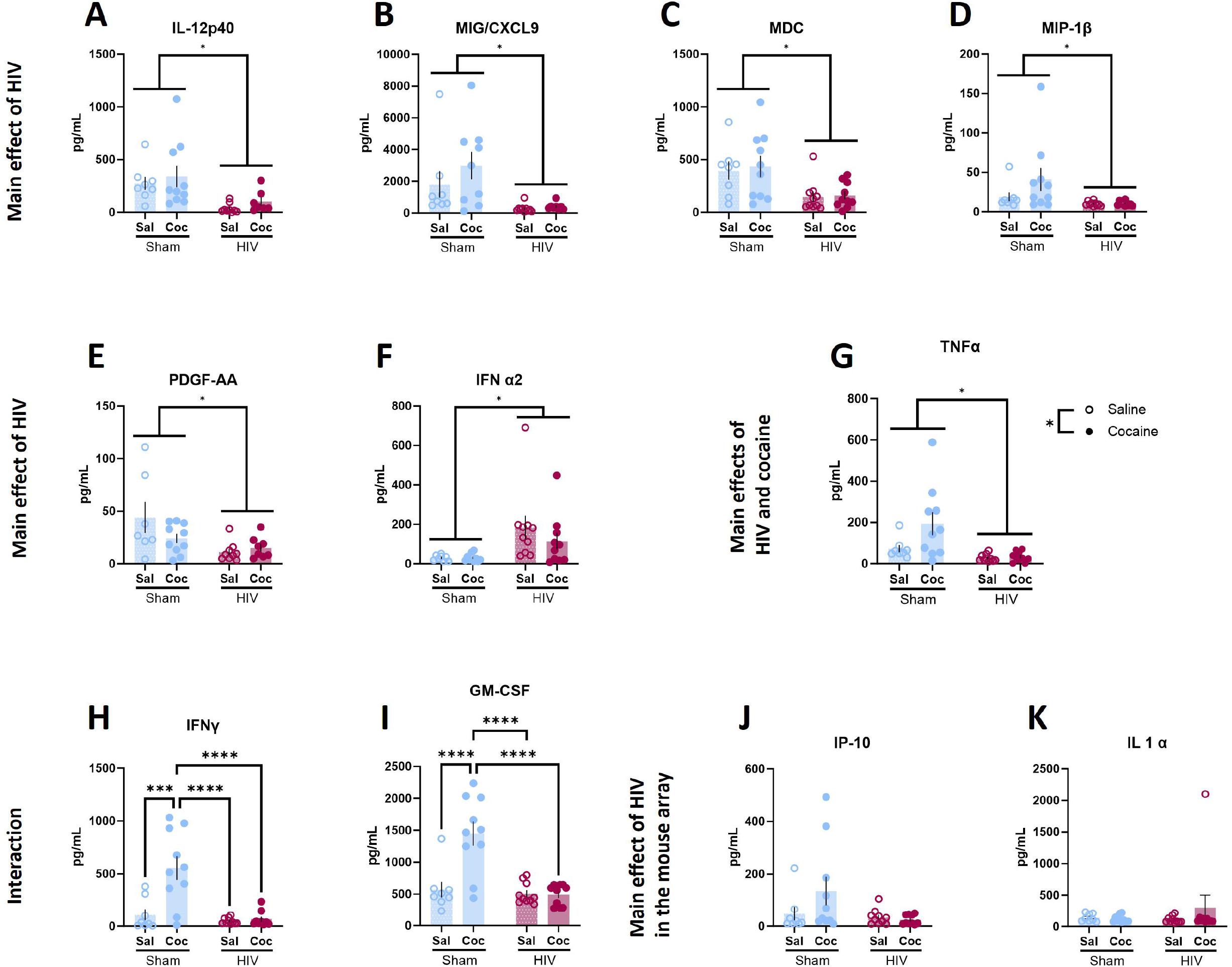
HIV and cocaine impacted human inflammatory factors in peripheral blood. (A, B, C, D, E, F) A main effect of HIV, but not cocaine, was observed in (A) IL-12p40, (B) MIG/CXCL9, (C) MDC, (D) MIP-1β, (E) PDGF-AA, and (F) IFN α2. (G) HIV and cocaine independently impacted TNFα expression, such that HIV-1 infected mice expressed lower TNFα, and cocaine exposure acted independently to increase expression. (H, I) HIV and cocaine interacted to determine (H) INFỿ and (I) GM-CSF levels. In sham mice, cocaine exposure increased INFỿ and GM-CSF, while this effect was absent in HIV-1 infected mice. (J, K) A subset of mouse targets was significantly altered by HIV-1 or cocaine, while no effects were observed in the HIV inflammatory factors. Analytes that differed in the mouse assay, but not the human assay were (J) IP-10, and (K) IL1α. *p < 0.05, ***p < 0.001, ****p < 0.0001. Data represent mean +/− SEM.

## Discussion

The current findings demonstrate that HIV-1 infection altered a subset of cocaine-related behaviors in mice, which was accompanied by peripheral inflammatory response and changes in brain microglia and astrocytes in neural substrates associated with cocaine reward and memory. HIV-1 infection did not impact acquisition of a cocaine CPP or reward seeking under conflict. In contrast, HIV-1 infected mice exhibited greater persistence of CPP under extinction conditions. Despite resistance to extinction of cocaine CPP, HIV-1 infected mice showed a similar magnitude of yohimbine-induced reinstatement to sham controls. These behavioral alterations were accompanied by changes in astrocyte immunoreactivity in the PFC such that HIV-1 infected mice showed greater GFAP staining in the IL and PL cortices. In contrast to HIV-1 effects on astrocytes, cocaine-induced alterations in microglia were observed.

HIV-1 infected humanized mice exhibited normal formation of a cocaine conditioned place preference, which assesses cocaine-reward associations with discrete contexts. This suggests that HIV-infected mice did not exhibit gross impairments in cocaine reward-related learning and did not have profound differences in cocaine reward compared to the sham-inoculated controls. Although initial CPP was similar, HIV-1 infected mice exhibited persistent expression of cocaine CPP across extinction sessions. Deficits in extinction of morphine CPP have been observed in the HIV-1 transgenic rat which expresses 7 of 9 HIV-1 viral proteins (Reid et al., 2001), though this model reflects consequences of protein exposure rather than infection. This may suggest generalized impairments in extinction across classes of rewards, though opioids likely interact with HIV to dysregulate cognitive function through discrete mechanisms from psychostimulants (Giacometti and Barker, 2019; Blackard and Sherman, 2021; Cirino and McLaughlin, 2021; Nash et al., 2021). The ability to extinguish reward seeking is dependent on the infralimbic cortex and its projections to the nucleus accumbens (Peters et al., 2008; LaLumiere et al., 2010, 2012). Deficits in extinction are particularly of note as our results identified increased GFAP immunoreactivity in the infralimbic and prelimbic cortices of HIV-infected mice, suggesting that prefrontal cortex function may be impaired. Indeed, others have found that astrocyte dysregulation in the medial prefrontal cortex is associated with extinction impairments in mice (Amaral et al., 2021; Taylor et al., 2021). This included reduced expression of the cystine-glutamate antiporter, xCT (Amaral et al., 2021). The cystine-glutamate antiporter is known to be upregulated by exposure to Tat, though it is not clear how this system is altered in the humanized mouse model. Both Tat and cocaine exposure independently alter astrocyte metabolism, resulting in reduced lactate transport to neurons, suggesting HIV-1 proteins may drive impairments in neuronal function indirectly through dysregulation of astrocytes (Natarajaseenivasan et al., 2018). We and others speculate that targeting astrocytes as regulators of dysregulated reward seeking behavior will be a promising therapeutic strategy across classes of addictive drugs (Scofield et al., 2015; Walker et al., 2020; Kim et al., 2022; Kruyer et al., 2022). The current findings suggest that astrocyte dysregulation is common across models of HIV, including infection and protein expression models, suggesting that astrocyte function and structure may further be promising for reversing HIV-associated changes in drug seeking behavior.

Yohimbine-induced reinstatement was used as a model of stress-induced relapse of drug seeking. Yohimbine produces physiological stress in humans and animals and is demonstrated to induce reinstatement of drug craving in clinical populations (Stine et al., 2002) and seeking in animal models (Anker and Carroll, 2010; Feltenstein et al., 2011; Brown et al., 2012). Despite overall extinction-resistance in HIV-1 infected mice, the magnitude of yohimbine-induced reinstatement was similar in the HIV-1 infected mice and sham-infected counterparts, suggesting that stress-induced relapse-related behavior is not facilitated in this model. Interestingly, in the Tat Tg models, Tat expression facilitated the reinstatement of ethanol CPP (McLaughlin et al., 2014), while the magnitude of cocaine-primed reinstatement was not altered in Tat Tg mice (Paris et al., 2014). This may suggest that reinstatement of cocaine seeking, a model of relapse-related behavior, is relatively spared from the effects of HIV infection or protein exposure in both models of pharmacological stressor-induced (yohimbine) and in drug-primed reinstatement.

Previous research into cocaine-related behavior in preclinical models of HIV has been largely limited to males. Our finding showing similar magnitude of CPP may result from differential effects of HIV-1 on cocaine preference in this model as compared to the use of Tat transgenic animals (Zhu et al., 2022), however, these previous findings were restricted to male mice. Substantial sex differences in cocaine-related behaviors have been reported, including development of cocaine CPP at lower doses in females (Zakharova et al., 2009) and reinstatement in females (Bobzean et al., 2010). While the current study did not detect sex differences, a majority of the subjects were female mice. Others have found that in female mice, Tat-facilitation of cocaine CPP was restricted to the diestrus phase (Paris, Fenwick et al., 2014). The diestrus phase is associated with elevated progesterone levels in mice (Walmer et al., 1992; Fata et al., 2001; Bastida et al., 2005; Maguire et al., 2005; Jaric et al., 2019) which we have shown to reduce stress-induced reinstatement (Feltenstein et al., 2011; Giacometti et al., 2022). As Tat induction can dysregulate neurosteroid synthesis (Paris et al., 2020; Qrareya et al., 2022) and treatment with progesterone or its metabolite allopregnanolone can attenuate both behavioral and neuropathological outcomes of Tat induction (Paris et al., 2016, 2020), neurosteroids are implicated in both the development and treatment of neurobehavioral impairments in HIV. Future studies into the relationship between HIV infection and behavioral outcomes should consider hormonal status as a potential mediator of these outcomes. As astrocytes in the prefrontal cortex – a key substrate of reinstatement – exhibited increased immunoreactivity in HIV-1 infected mice in this model, and they further act as key regulators of steroid signaling in the brain (Micevych and Sinchak, 2008; Azcoitia et al., 2010), this further underscores the importance of considering circulating hormones and non-neuronal contributors in neuroHIV outcomes.

In contrast to prefrontal astrocyte changes induced by HIV-1 infection, alterations in microglia populations induced by cocaine exposure were observed in the basolateral amygdala and the CA1 subfield of the hippocampus. While cocaine exposure increased overall Iba1 staining in the basolateral amygdala - indicative of cocaine-induced morphology changes consistent with activated microglia in HIV-1 infected and uninfected mice, cocaine only increased microglia cell counts in the hippocampus of HIV-1 infected mice. This represents a unique vulnerability of the hippocampus to microglial dysregulation by cocaine in the context of HIV-1. The hippocampus is essential for normal learning and memory processes, including extinction of cocaine conditioned place preference (Hitchcock and Lattal, 2018), which was impaired in HIV-1 infected mice. These findings suggest a different profile of hippocampal microglia response to systemic HIV-1 infection when compared to local infusion of HIV-1 proteins to the hippocampus, which induces microglial activation (Hill et al., 2019). This may reflect higher exposure to HIV-1 proteins resulting from this local infusion to the tissue. Hippocampal vulnerability in the EcoHIV model of systemic infection has also been reported (Kelschenbach et al., 2019; Kim et al., 2019), which is accompanied by altered hippocampus synaptodendritic integrity. Hippocampus network function has also been shown to be dysregulated by both exposure to HIV-1 Tat and cocaine (Ahooyi et al., 2018). Notably, cocaine and Tat-induced deficits in hippocampus network activity were attenuated in the presence of astrocytes. While we did not observe changes in GFAP immunoreactivity in the hippocampus of HIV-1 infected mice, these blunt measures may miss changes in astrocytic synaptic coverage or function which could contribute to changes in hippocampal synaptodendritic integrity or plasticity when exposed to cocaine or elevated dopamine levels, resulting in aberrant hippocampal function. Importantly, although the morphology of Iba1-labeled cells was consistent with microglia, Iba1 can also label macrophages. As HIV infection may increase peripheral immune cell migration into the central nervous system, it is possible that macrophage populations contribute to observed effects of HIV on Iba1 expressing populations. Because this model includes humanization of the peripheral immune system, changes in the CNS to astrocyte and microglia populations as a result of HIV-1 infection likely in part reflect outcomes of peripheral infection and underscore the effects of inflammation on the brain and behavior.

These neural changes were accompanied by inflammatory changes in peripheral blood following behavioral testing. To investigate changes in both humanized and mouse-derived systems, we assessed changes in both human (derived from humanized immune cells) and mouse (derived from host mouse cells) cytokines and chemokines. In both the mouse and human arrays, a majority of the observed changes were driven by HIV-1 infection. For mouse targets, this included downregulations of IL-1α and the granulocyte colony stimulating factor (G-CSF) and upregulations in RANTES/CCL5 and IP-10/CXCL10, which have been previously implicated in multiple models of HIV infection and in clinical populations, including in co-occurring cocaine use/expose and infection (Nair et al., 2000; Parikh et al., 2014; Nayak et al., 2020). Elevated G-CSF has also been reported to increase dopamine release in response to cocaine (Brady et al., 2019, 2021), though as with other considerations this may be mediated by hormonal status in female mice (Brady et al., 2019). Though we did not observe main effects of cocaine itself on mouse G-CSF levels, the cocaine exposure paradigms in the current study were at lower doses (10 mg/kg vs 20 mg/kg) and fewer exposures (3 vs 7 injections), suggesting that a more protracted cocaine exposure regimen may be necessary to observe cocaine effects or cocaine and HIV interactive effects on G-CSF expression.

Human granulocyte-macrophage colony stimulating factor (GM-CSF) levels were increased by cocaine exposure in uninfected mice, while this effect was blunted in mice infected with HIV-1. This aligns with previous findings that in a cocaine sensitization model, repeated cocaine exposure increased interferon gamma levels in immunointact mice (Kubera et al., 2004). Further, cocaine-induced increases in GM-CSF have been observed in other models of experimenter administered cocaine (Calipari et al., 2018) under conditions that similarly induced increases in G-CSF. In PLWH who use drugs, GM-CSF levels were significantly associated with historical cocaine use (Parikh et al), identifying dysregulation of GM-CSF as a common target in both preclinical and clinical data sets. Notably, human targets that were modulated by cocaine were generally pro-inflammatory, with cocaine-induced increases in sham mice in interferon gamma as well as GM-CSF and increases in TNF in both sham and HIV-infected mice. Consistent with findings from the literature, this may implicate cocaine in producing a pro-inflammatory state and generalized impairment in immune function. The current findings reflect repeated, but not chronic, cocaine exposure, and thus it will be critical to expand these results to models of chronic drug exposure and dependence. To our knowledge, a mechanistic role of GM-CSF and interferon gamma have not been assessed in the regulation of cocaine-related behavior either systemically or in reward-related neural structures, suggesting a potential avenue for reversal of HIV-1 associated alterations in cocaine reward-related behaviors.

The current study provides additional understanding of the interaction between HIV infection and cocaine exposure to alter the immune landscape and promote continued drug seeking. This area of investigation has faced challenges due to difficulties in modeling HIV infection in preclinical models. This study used a model in which mice are humanized by a process in which the mouse’s immune cells are irradiated at birth, and human HPSCs are engrafted. The humanized mouse model thus offers insight into a number of unique questions, including how HIV-1 infection *per se* as opposed to protein exposure alone, is impacting central and peripheral inflammation and impacting behavior. These findings have complemented findings from other models, including the ecoHIV model and Tat transgenic model, with several converging areas – such as extinction learning and neuroimmune function –, which we expect to be particularly robust targets for future investigation. Limitations of the current model included behavioral abnormalities such as high rates of stereotypy of mice and differences in overall brain morphology, which prevented automation of analyses and will require robust calibration for future work manipulating discrete neural circuits as neuroanatomical differences between the humanized mice and commonly used inbred mouse lines were the norm.

Together, findings in these studies demonstrate that, in mice with humanized immune systems, HIV-1 infection is associated with deficits in extinction of cocaine conditioned place preference. This is associated further with astrocyte and microglia alterations in key substrates of reward extinction, including subregions of the medial prefrontal cortices and the hippocampus. Further, systemic inflammatory alterations resulted from HIV-1 infection and cocaine exposure, both independently and in tandem, which identifies additional targets of investigation for peripheral immune consequences within the central nervous system that may contribute to deficits in the regulation of cocaine seeking behavior in the context of HIV-1 infection.

## Acknowledgements

This research was supported by NIH awards DP2DA051907 (JMB), R03DA047919 (JMB), 1R01DA054535-01 (SG) and 1R33DA041018-01 (SG), and a pilot award from The Comprehensive Neuro-AIDS Center Grant P30MH092177-9 (JMB and LLG).

## Figure Legends

**Supplemental Figure 1.** IBA1 cells per mm2 in the (A) CA2, (B) CA3, and (C) DG of the hippocampus. Percent area of IBA1 in the (D) CA2, (E) CA3, and (F) DG of the hippocampus.

