## Supplementary figures and images for "Impaired extinction of cocaine seeking in HIV-infected mice is accompanied by peripheral and central immune dysregulation"

### Supplemental Figure 1

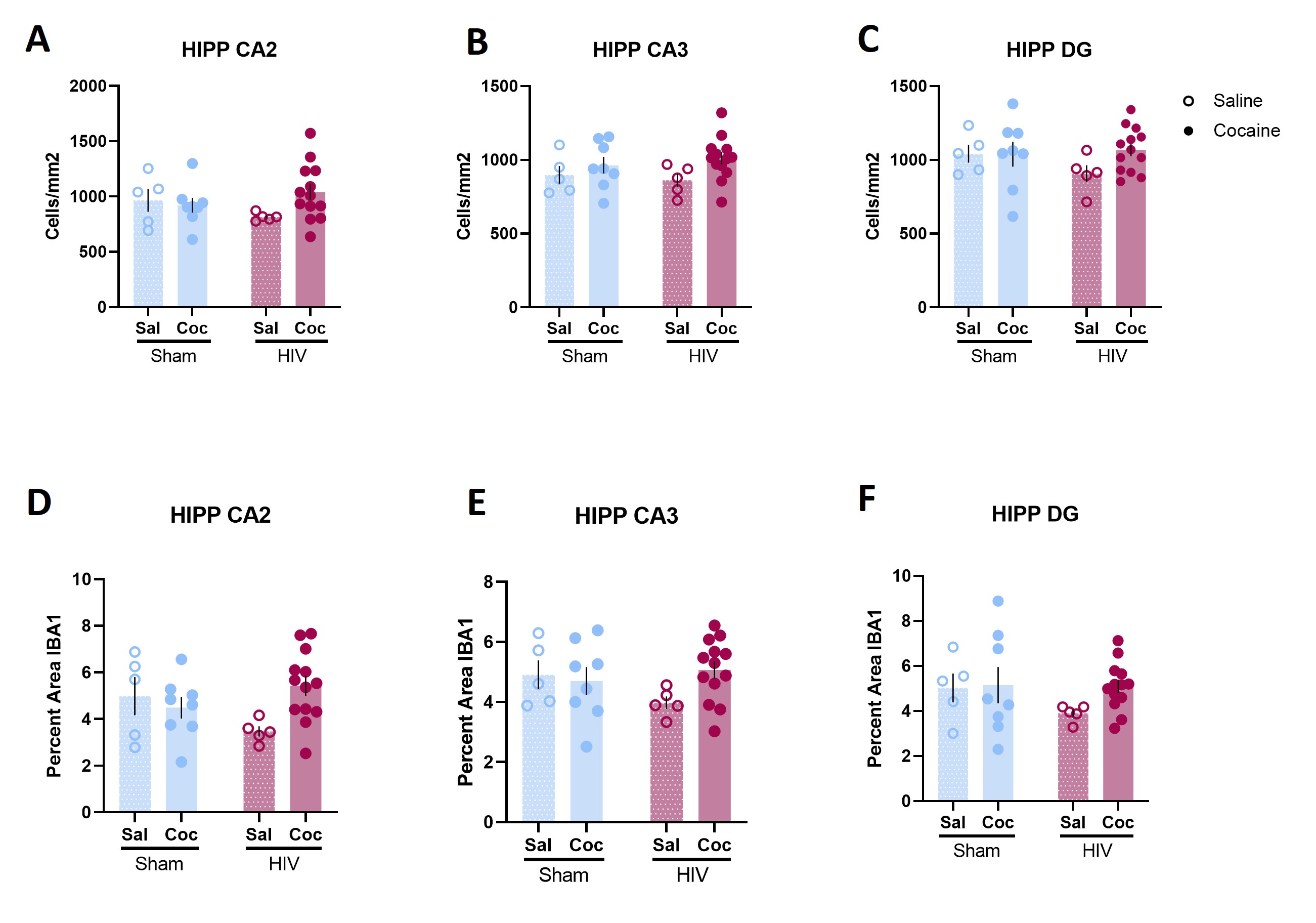
